# Functional immune state classification of unlabeled live human monocytes using holotomography and machine learning

**DOI:** 10.1101/2023.09.12.557503

**Authors:** Mahn Jae Lee, Geon Kim, Moo Sung Lee, Jeong Won Shin, Jung Ho Lee, Dong Hun Ryu, Young Seo Kim, YoonJae Chung, Kyuseok Kim, YongKeun Park

## Abstract

Precise evaluation of immune status is critical for managing diseases such as sepsis, in which the immune system transitions between hyper-inflammatory and immune-suppressed states. However, current biomarkers are limited by low specificity and time-consuming protocols. Here, we present a label-free, imaging-based framework for single-cell immune profiling of human monocytes using three-dimensional holotomography (HT) and deep learning. HT captures subcellular refractive index (RI) distributions of live, unlabeled cells, enabling quantitative extraction of morphological and biophysical features. Using an *in vitro* LPS-based model, we classified three functional immune states—control, hyper-inflammation, and immune suppression—based on 4,059 holotomograms from 11 donors. Immune state transitions were associated with significant changes in cell volume, surface area, RI variability, and the abundance of lipid droplets. A 3D convolutional neural network trained on HT images achieved 83.7% accuracy for single-cell predictions, increasing to 99.9% with ensemble averaging. This study establishes HT as a scalable, label-free platform for real-time immune monitoring and introduces subcellular RI features as robust correlates of immune dysregulation.

## Introduction

Classical immunology drew heavily on cell shape—e.g., the lobulated neutrophil nucleus or the dendrite-studded dendritic cell—as a first discriminator of lineage^1^. Although modern “omics” technologies have largely replaced morphology in immunophenotyping, cell shape remains a rapid, information -dense, and increasingly quantifiable read-out of immune function^2^. Systematic mapping of morphological determinants has the potential to refine our understanding of immune regulation, generate orthogonal biomarkers, and inspire morphology-guided therapeutic strategies.

Recent advances support the view that immune cell morphology reflects not just identity but also functional state. Distinct morphological features—including nuclear geometry, cytoplasmic asymmetry, organelle positioning, and membrane dynamics—have been correlated with immune activation, polarization, exhaustion, and effector function across both innate and adaptive immune cells. For instance, neutrophil nuclear delobulation presages NETosis^3^, elongated macrophage shape promotes M2 polarization ^4^, and fragmented immunological synapses signal T-cell exhaustion^5,6^. These morphological signatures are not merely descriptive but reflect underlying transcriptional and metabolic programs. Emerging label-free 3D imaging techniques such as holotomography (HT), when coupled with deep-learning morphometrics, enable high-throughput, quantitative profiling of these shape-based biomarkers at single-cell resolution^7^. Such approaches are transforming morphology into a real-time, scalable biomarker for immune monitoring and precision diagnostics.

These capabilities are particularly relevant in clinical settings like sepsis, where the immune system exhibits a dynamic shift from hyper-activation to suppression^8^. Sepsis is a life-threatening condition resulting from a dysregulated immune response to infection ^9^, initially marked by a surge of inflammatory cytokines and progressing toward an immune-suppressed state. This immunological trajectory is associated with poor outcomes, including septic shock, secondary infections, and increased mortality ^8,10^. Immunostimulatory agents such as granulocyte-macrophage colony-stimulating factor, interferon-gamma, and immune checkpoint inhibitors have been proposed to reverse immune paralysis ^11-13^. However, timely and accurate assessment of immune status remains a major clinical challenge.

While current biomarkers such as C-reactive protein and procalcitonin are routinely used in sepsis management, their limited specificity and high inter-individual variability reduce diagnostic reliability ^14,15^. One promising marker is the expression of HLA-DR on monocytes (mHLA-DR), which correlates with immune suppression and prognosis ^16-19^. However, mHLA-DR measurement requires laborious, multi-step labeling and yields only semi-quantitative data, hindering real-time clinical application.

In this study, we apply HT and deep learning to classify the immune status of individual monocytes by capturing intracellular alterations associated with immune dysregulation. Unlike previous approaches focused on cell identity classification, our method predicts functional immune states, enabling rapid, label-free immune assessment. HT reconstructs the RI distribution of live, unlabeled monocytes at subcellular resolution without exogenous labeling. From these 3D RI tomograms, we extracted quantitative morphological and biophysical features—termed HT features—across three defined immune conditions: control, hyper-inflammation, and immune suppression. These features exhibited significant changes with immune state progression, including alterations in intracellular mean RI. A deep learning classifier trained on these features achieved 83.7% accuracy for single-cell predictions, which improved to 99.9% when aggregating results from six monocytes. This approach provides a robust, real-time platform for functional immune profiling, with potential applications in sepsis management and broader immune monitoring.

## Results

### Holotomograms of monocytes in different immune statuses

To model immune dysregulation in sepsis, we applied lipopolysaccharide (LPS) stimulation to induce distinct immune states in monocytes. Although based on an *in vitro* system, these conditions are well-established functional analogues of clinical immune activation and suppression, as supported by previous studies ^20,21^. In this study, we adopt these experimental states as proxies for clinically relevant immune phenotypes. Monocytes were enriched from peripheral blood mononuclear cells (PBMCs) using magnetic-activated cell sorting (MACS) and subsequently differentiated into three groups—control, hyper-inflammation, and immune suppression—following the established protocol (Fig. 1a; see Methods for details). Cell viability was consistently greater than 95% across all conditions.

**Fig. 1.**
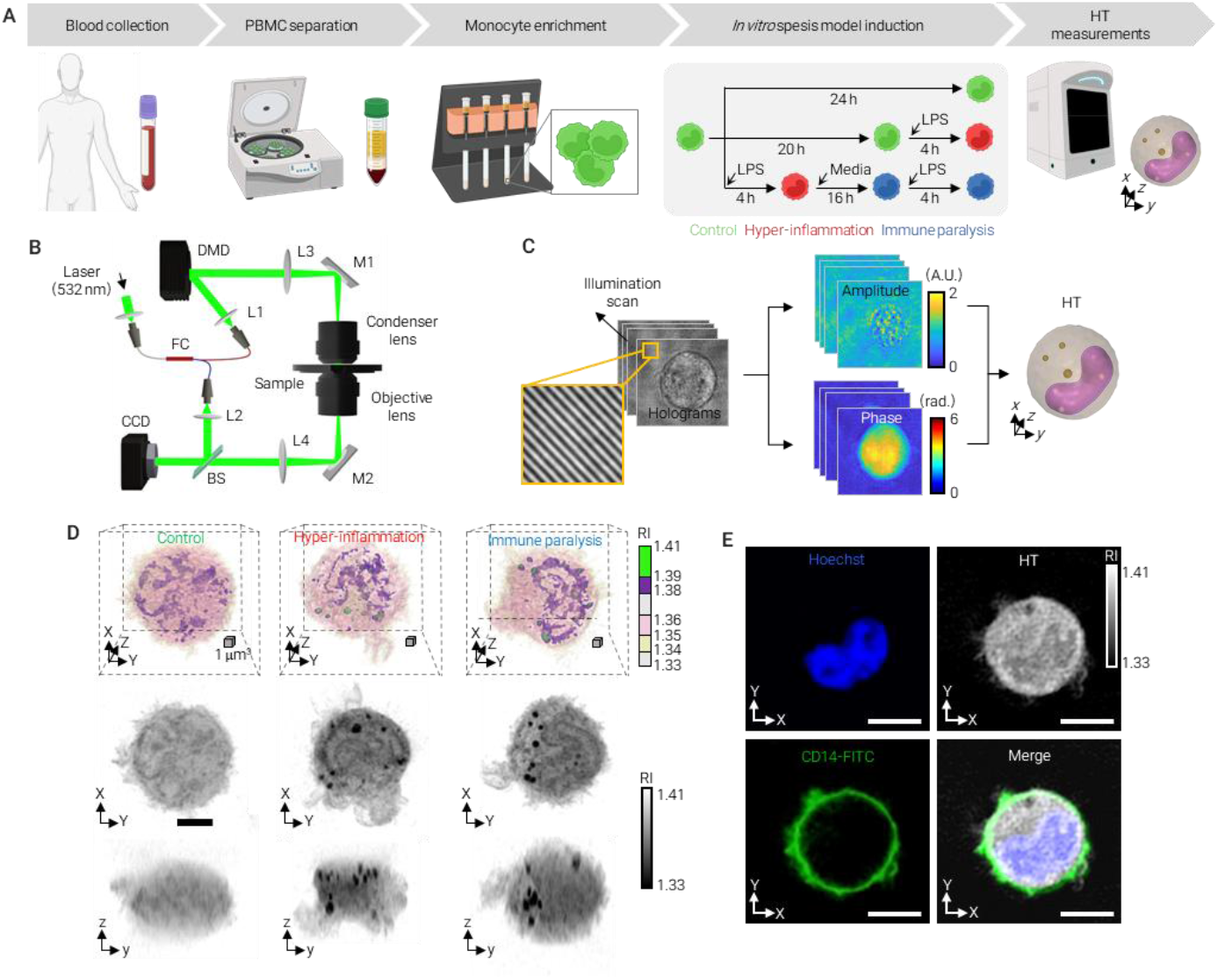
Experimental workflow and representative holotomograms of monocytes. a, Experimental scheme. Monocytes were isolated from PBMCs using MACS and cultured under control (green), hyp er-inflammatory (red), or immune-suppressed (blue) conditions based on an in vitro LPS model. b, Optical layout of the holotomography system using a Mach–Zehnder interferometer. c, RI tomogram reconstruction from multiple 2D holograms acquired at different illumination angles. d, Representative 3D RI images of monocytes in different immune states. Left: 3D renderings. Right: maxima l intensity projections (XY, YZ, XZ). Scale bars = 5 μm. e, Correlative fluorescence images showing CD14 (green, FITC) and nucleus (blue, Hoechst) overlaid with RI map. Scale bars = 5 μm.

To acquire three-dimensional RI distributions of individual unlabeled monocytes, we employed a commercial HT system (HT-2H, Tomocube Inc., Republic of Korea). This system utilizes a Mach–Zehnder interferometric setup with a coherent 532 nm laser and a digital micromirror device (DMD) to deliver multi-angle illumination^22-24^. A total of 49 off-axis holograms were captured per cell under varied illumination angles and computationally reconstructed into volumetric RI tomograms by solving the inverse Helmholtz equation ^25-28^. The acquisition time per cell was 0.1 seconds, with minimal light exposure (<10 μW at the sample plane), and the system achieved a theoretical resolution of 110 nm laterally and 330 nm axially. This approach enabled label-free, high-resolution visualization of subcellular structures such as the nucleus, nucleoli, and cytoplasmic granules, laying the foundation for subsequent quantitative feature extraction and immune state classification.

The reconstructed RI tomograms clearly revealed subcellular features of individual monocytes, including the nucleus, nucleoli, and cytoplasmic granules (Fig. 1d). Morphological differences between immune states were readily apparent. Monocytes in hyper-inflammatory and immune-suppressed conditions exhibited altered cytoplasmic RI distributions, as well as increased numbers of discrete high-RI granules, compared to control cells. These high-RI domains, later identified as lipid droplets, appeared more frequently and prominently in immune-dysregulated states. In addition to cytoplasmic features, variations in nuclear size, membrane curvature, and internal RI heterogeneity were also observed across immune conditions. To confirm cell identity and validate segmentation boundaries, we performed correlative fluorescence imaging with FITC-conjugated anti-CD14 antibodies and Hoechst nuclear staining (Fig. 1e). All imaged cells exhibited strong CD14 expression and a single, clearly delineated nucleus, confirming their monocyte identity and morphological integrity. These high-resolution, label-free 3D holotomograms provided the structural foundation for subsequent quantitative feature extraction and immune state classification.

### HT data collection and feature extraction

To quantitatively assess morphological and biophysical features associated with immune status, we analyzed a total of 4,059 monocyte holotomograms collected from 11 healthy donors. For each donor, PBMCs were isolated and subjected to LPS stimulation to induce three immune states—control, hyper-inflammation, and immune suppression—following a standardized in vitro protocol (see Methods).

Holotomograms were segmented using the Otsu thresholding method to define the cell boundary in three dimensions (Fig. 2a)^29-31^. Within each segmented cell volume, we computed geometric parameters including cell volume, surface area, and aspect ratio. In addition, RI-based features such as mean RI, standard deviation of RI, and estimated dry mass were extracted. The measured RI values ranged from 1.33 to 1.41, consistent with known values for biological cells (Fig. 2b)^32,33^.

**Fig. 2.**
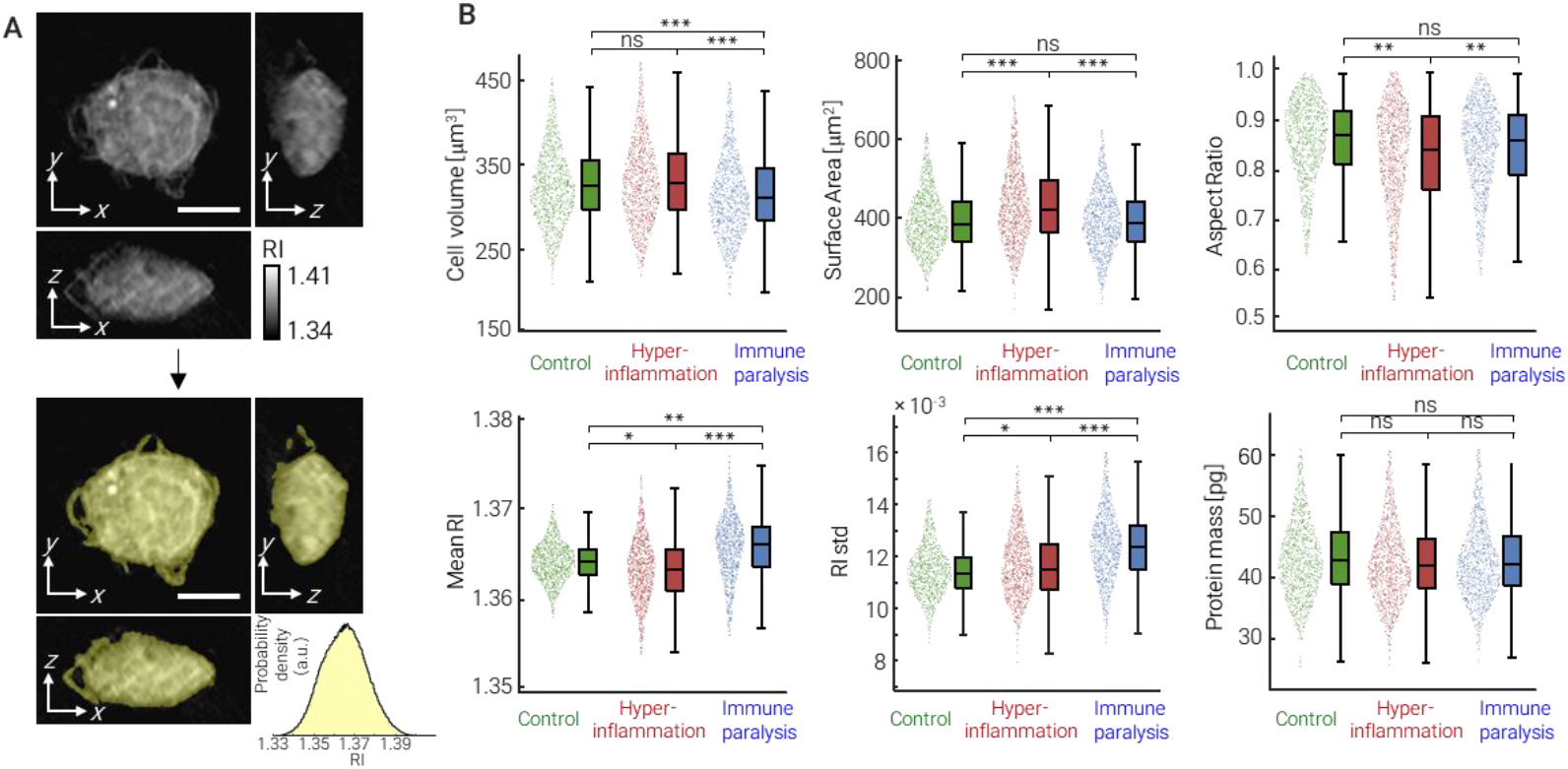
Quantitative analysis of morphological and biophysical features of individual monocytes across immune states. a, Representative holotomographic images of a monocyte and its 3D segmentation using the Otsu method. RI maps and segmented boundaries are shown. Scale bars = 5 μm. b, Refractive index (RI) distribution of the segmented monocyte shown in (a), ranging from 1.33 to 1.41. c, Comparison of extracted features among control (green), hyper-inflammatory (red), and immune-suppressed (blue) monocytes: cell volume, surface area, aspect ratio, mean RI, RI standard deviation, and estimated dry mass. Violin plots show the distribution of values; boxplots indicate medians and interquartile ranges. Statistical significance was assessed using a two-sided unpaired Student’s t-test. ns, not significant; *P < 0.05; **P < 0.01; ***P < 0.001.

Given that RI correlates linearly with protein concentration in aqueous media ^34,35^, we calculated the intracellular dry mass by applying a RI increment (α = 0.185 mL/g) to the voxel-wise RI distribution. This enabled label-free quantification of total protein content for each cell. Together, these features provided a multidimensional descriptor of monocyte morphology and composition, which served as input for downstream statistical analysis and machine learning-based immune state classification (Fig. 2c).

### Quantitative morphological analysis of individual monocytes in various immune states

Quantitative analysis of holotomograms revealed distinct morphological and biophysical profiles of monocytes across immune states. Cells under immune suppression exhibited a significant reduction in volume (306.4 ± 51.6 μm^3^) compared to control (318.2 ± 49.0 μm^3^) and hyper-inflammatory monocytes (329.79 ± 55.9 μm^3^). In contrast, surface area was highest in the hyper-inflammation group (414.6 ± 79.8 μ m^2^), exceeding that of both control (385.1 ± 67.4 μm^2^) and immune-suppressed cells (384.8 ± 72.6 μm^2^), suggesting increased membrane ruffling and spreading in activated states.

Aspect ratio, an indicator of cellular symmetry, was reduced in hyper-inflammatory monocytes (0.819 ± 0.103), reflecting more irregular shapes, while control and immune-suppressed cells exhibited relatively more symmetric morphologies (0.852 and 0.834, respectively).

RI analysis showed a subtle but consistent trend: mean RI was lowest in hyper-inflammatory cells (1.360 ± 0.003), intermediate in control (1.361 ± 0.002), and highest in immune-suppressed monocytes (1.362 ± 0.003), indicating shifts in intracellular composition. Despite these RI variations, estimated total dry mass remained approximately constant (∼45 pg), suggesting that functional immune transitions are accompanied by structural reorganization rather than net changes in protein content.

### Identification of High-RI Granules as Lipid Droplets in Monocytes under Septic Conditions

HT revealed an increased number of intracellular granules with high RI values in monocytes exposed to septic conditions, compared to controls (Fig. 3a). Based on previous reports, we hypothesized that these high -RI structures correspond to lipid droplets (LDs)^27,35^. To test this, we performed BODIPY staining, a fluorescent marker for neutral lipids, and compared the labeled regions with HT-segmented granules across varying RI thresholds using maximum intensity projections (Fig. 3b). The best overlap was obs erved at RI > 1.39, consistent with prior studies, supporting the identification of high-RI granules as LDs (Fig. 3c)^27^.

**Fig. 3.**
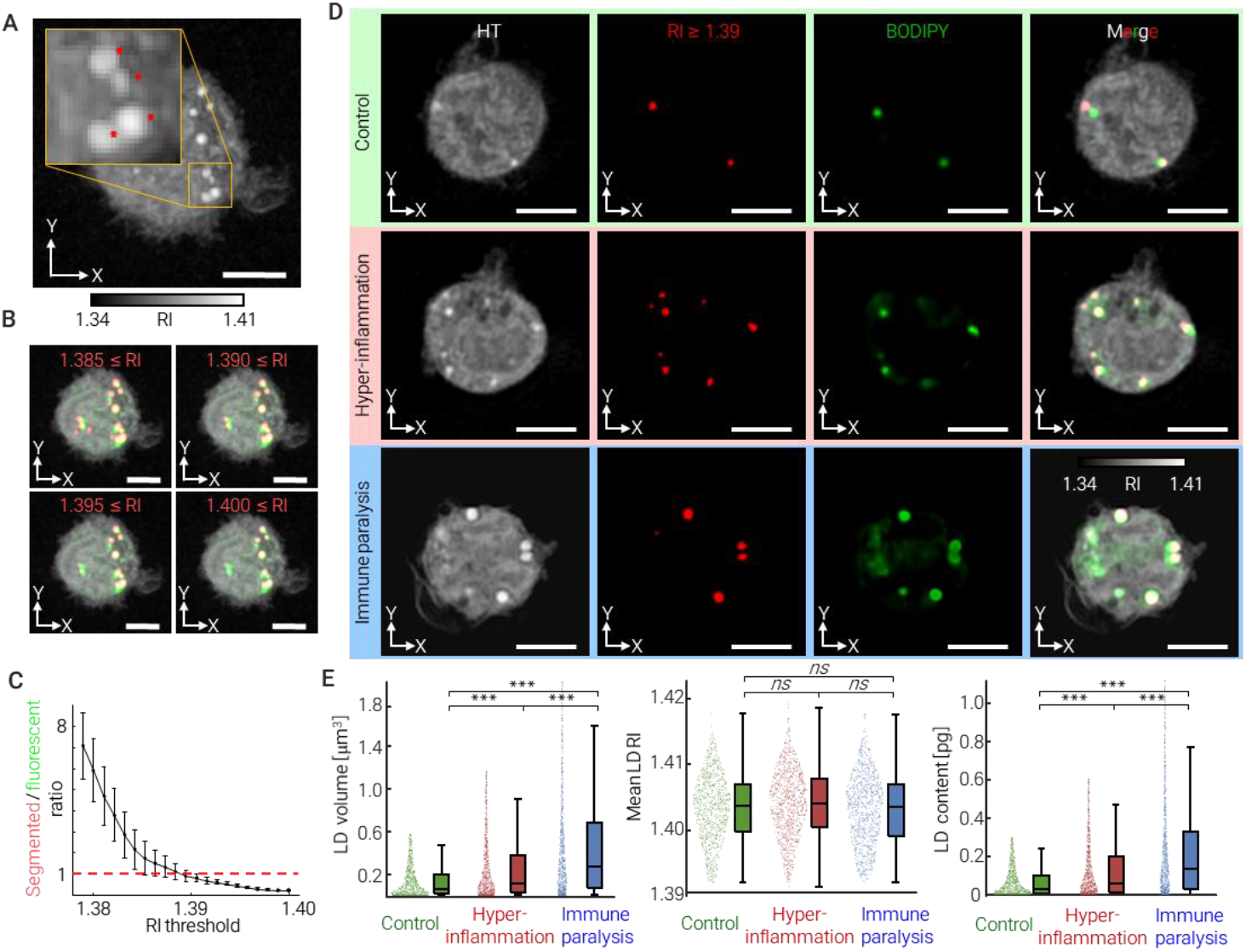
Quantitative identification and analysis of intracellular LDs in monocytes. a, Representative RI tomogram showing intracellular high-RI granules (red asterisks), magnified in the inset. Scale bar = 5 μm. b, RI-based segmentation of granules using threshold values from 1.385 to 1.400, overlaid with BODIPY fluorescence (green). Scale bars = 5 μm. c, Area overlap ratio between segmented high-RI regions and fluorescently labeled LDs as a function of RI threshold. d, Correlative imaging of monocytes under different immune states (control, hyper-inflammation, immune suppression) showing RI maps (gray), segmented LDs (red), BODIPY signals (green), and merged views. Scale bars = 5 μm. e, Statistical comparison of LD volume, mean RI, and estimated LD mass across immune states. Violin plots show distributions; boxplots indicate medians and interquartile ranges. Statistical significance was assessed by two-sided unpaired Student’s t-test. ns, not significant; *P < 0.05; **P < 0.01; ***P < 0.001.

Using this threshold, we segmented and quantified LDs in monocytes across immune conditions (Fig. 3d). LD volume increased progressively with immune activation and suppression: 0.62 ± 0.59 μm^3^ in control, 1.14 ± 1.13 μm^3^ in hyper-inflammatory, and 1.99 ± 1.88 μm^3^ in immune-suppressed cells. Despite these volume changes, the average RI of LDs remained consistent (∼1.396), indicating stable lipid concentration. Applying a known RI increment of 0.135 ml/g, we estimated LD mass per cell, which also increased with immune dysregulation: 0.28 ± 0.27 pg (control), 0.52 ± 0.53 pg (hyper-inflammation), and 0.88 ± 0.83 pg (immune suppression) (Fig. 3e).

### The Correlation Between Immune Status and HT Features

To capture subtle differences in intracellular RI, we subdivided each monocyte holotomogram into six RI intervals (I–VI) with a resolution of 0.005 (Fig. 4a). From each subrange, we extracted spatial and morphological features—collectively referred to as HT features—based on voxel distribution and distance from the cell centroid (Fig. 4b, Supplementaly Information - Table 1).

**Fig. 4.**
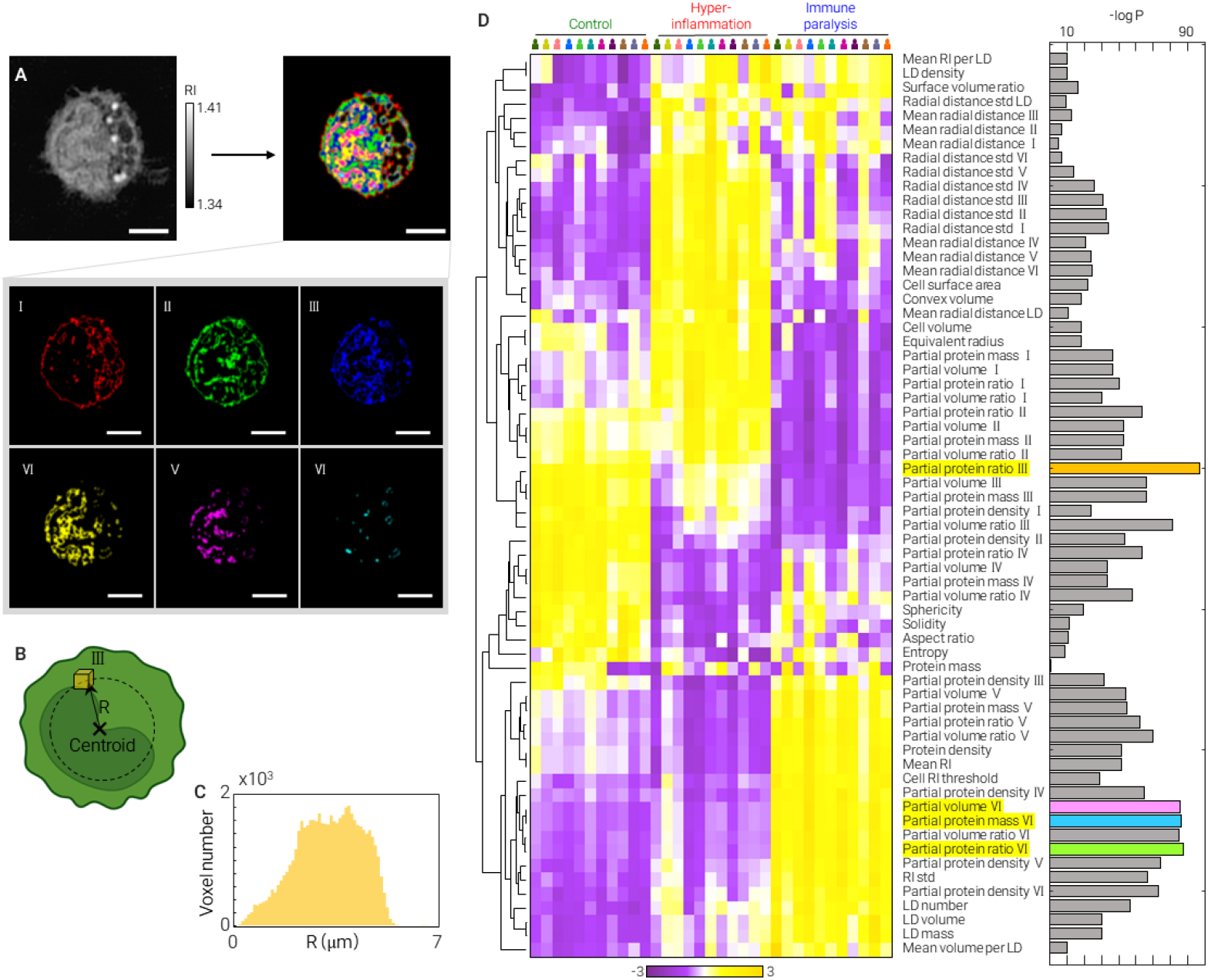
Morphological and refractive index-based feature mapping of monocytes across immune states. a, Segmentation scheme dividing monocyte holotomograms into six RI subranges for partial feature extraction. Scale bar = 5 μm. b, Schematic illustration of partial HT feature calculation based on voxel distance from the centroid, shown for RI subrange III (1.365–1.370). c, Z-score heatmap of 64 HT features across individual donors and immune states. Rows represent features; columns represent averaged donor samples. Dendrogram reflects inter-feature correlation. d, Multivariate analysis of variance (MANOVA) ranking feature importance for immune status discrimination. The top four most distinguishable features are color-highlighted.

We analyzed 64 HT features across immune states using feature-based machine learning. Z-score normalization and heatmap visualization revealed distinct feature patterns associated with each immune condition (Fig. 4c). Notably, LD-related features were elevated in both hyper-inflammatory and immune-suppressed monocytes, consistent with previous findings (Fig. 3e). Additionally, monocytes in the hyper-inflammatory state showed greater RI variability across all subranges, indicating increased intracellular heterogeneity.

To identify features most predictive of immune status, we applied multivariate analysis of variance (MANOVA) to rank all 64 features by statistical significance (Fig. 4d). The partial protein ratio in RI subrange III (1.365–1.370) emerged as the most discriminative feature, whereas total protein mass showed minimal variation across conditions (see Fig. 2c), suggesting it is not a reliable immune status marker.

### A Multidimensional Analysis of Immune Status in HT Feature Space

To visualize the distribution of HT features and their relationship to immune status, we projected 64 - dimensional feature vectors of individual monocytes into a two-dimensional space using Uniform Manifold Approximation and Projection (UMAP) (Fig. 5a). Cells with the same immune status formed distinct clusters. Gaussian kernel density estimation was applied to each group, and 1σ boundaries from the centroid were drawn to define characteristic regions in feature space (Fig. 5b)^36^. Within each gated region, the majority of monocytes belonged to the corresponding immune class (Fig. 5c), supporting the utility of HT features in distinguishing immune states.

**Fig. 5.**
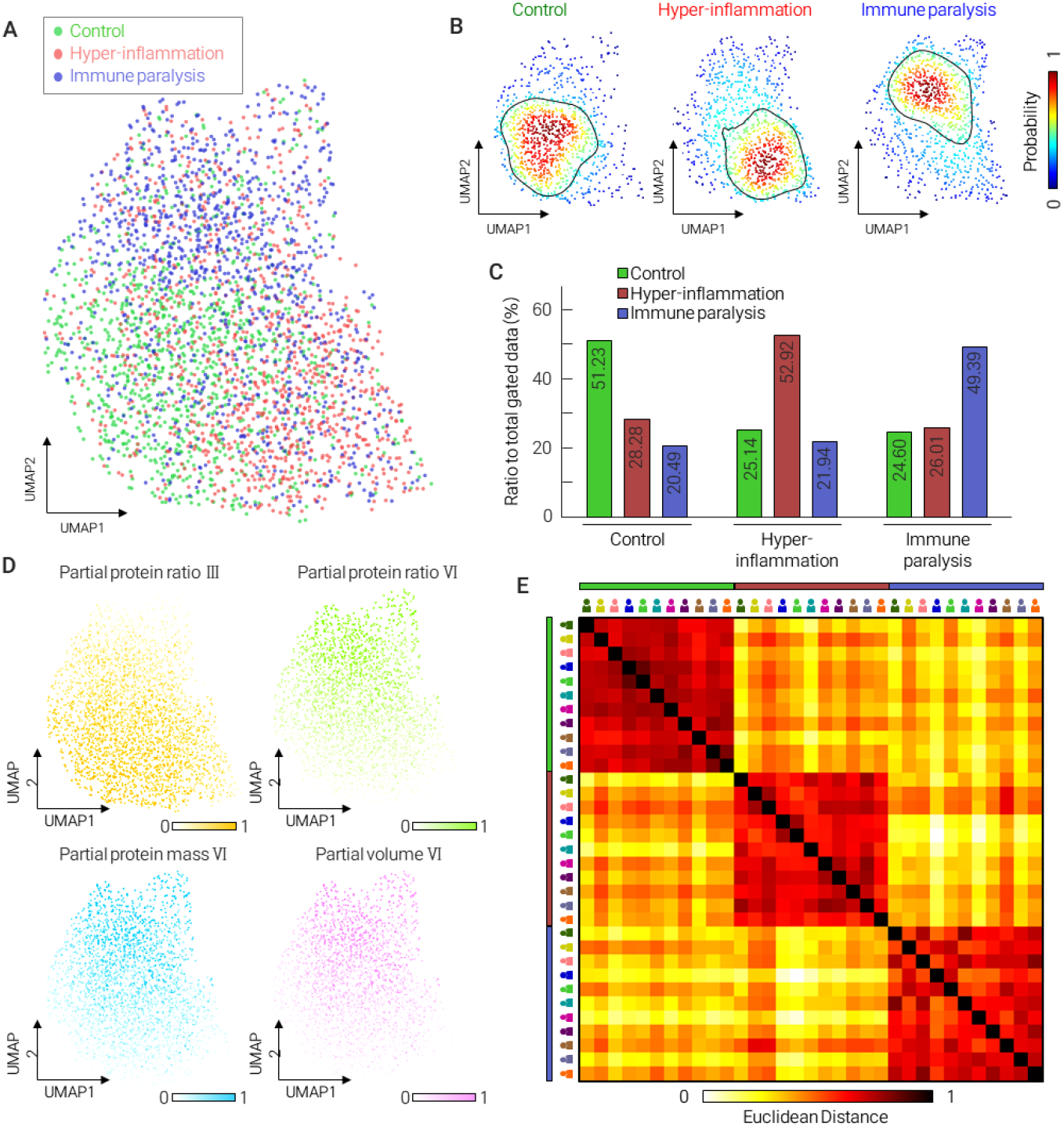
Immune status classification of monocytes using HT features and machine learning. a, UMAP projection of 64 HT features from individual monocytes. Each point represents a single cell, color-coded by immune status (green: control, red: hyper-inflammation, blue: immune suppression). b, Gaussian kernel density estimation for each immune state in UMAP space. One standard deviation (1σ) boundary from each centroid is outlined. c, Proportion of monocytes from each immune status within the 1σ gated regions shown in (b). d, UMAP heatmaps showing the spatial distribution of four representative HT features: partial protein ratio III, partial protein ratio VI, partial protein mass VI, and partial volume VI. e, Normalized Euclidean distance heatmap of HT feature profiles across individual donors and immune statuses. Donor groups are annotated by colored bars.

To identify features driving the separation, we mapped the top four most discriminative features onto the UMAP space (Fig. 5d). The partial protein ratio III was enriched in control cells, while partial volume VI, partial protein ratio VI, and partial protein mass VI were predominant in the immune suppression group. These spatial distributions coincided with the density clusters observed in Fig. 5b, suggesting that specific RI-based features underlie immune state–dependent phenotypes.

To evaluate inter-donor consistency, we computed the Euclidean distances between averaged HT feature profiles of monocytes grouped by donor and immune status. As shown in the heatmap (Fig. 5e), distances between different immune statuses were significantly greater than those between donors within the same immune group. This indicates that immune status–dependent differences in HT features outweigh donor-specific variability, supporting the robustness of our method and its potential as a generalizable immune profiling tool.

### Immune Status Prediction Using Deep Learning Classification

We developed a deep learning classifier to predict immune status directly from monocyte holotomograms, leveraging the distinct HT feature profiles observed across immune states. The model architecture was based on a 3D DenseNet framework comprising four dense blocks with 12, 24, 64, and 64 convolutional layers, respectively (Fig. 6a). This structure enables multi-scale feature extraction and efficient reuse of spatial information, well suited for capturing subtle variations in refractive index distributions.

**Fig. 6.**
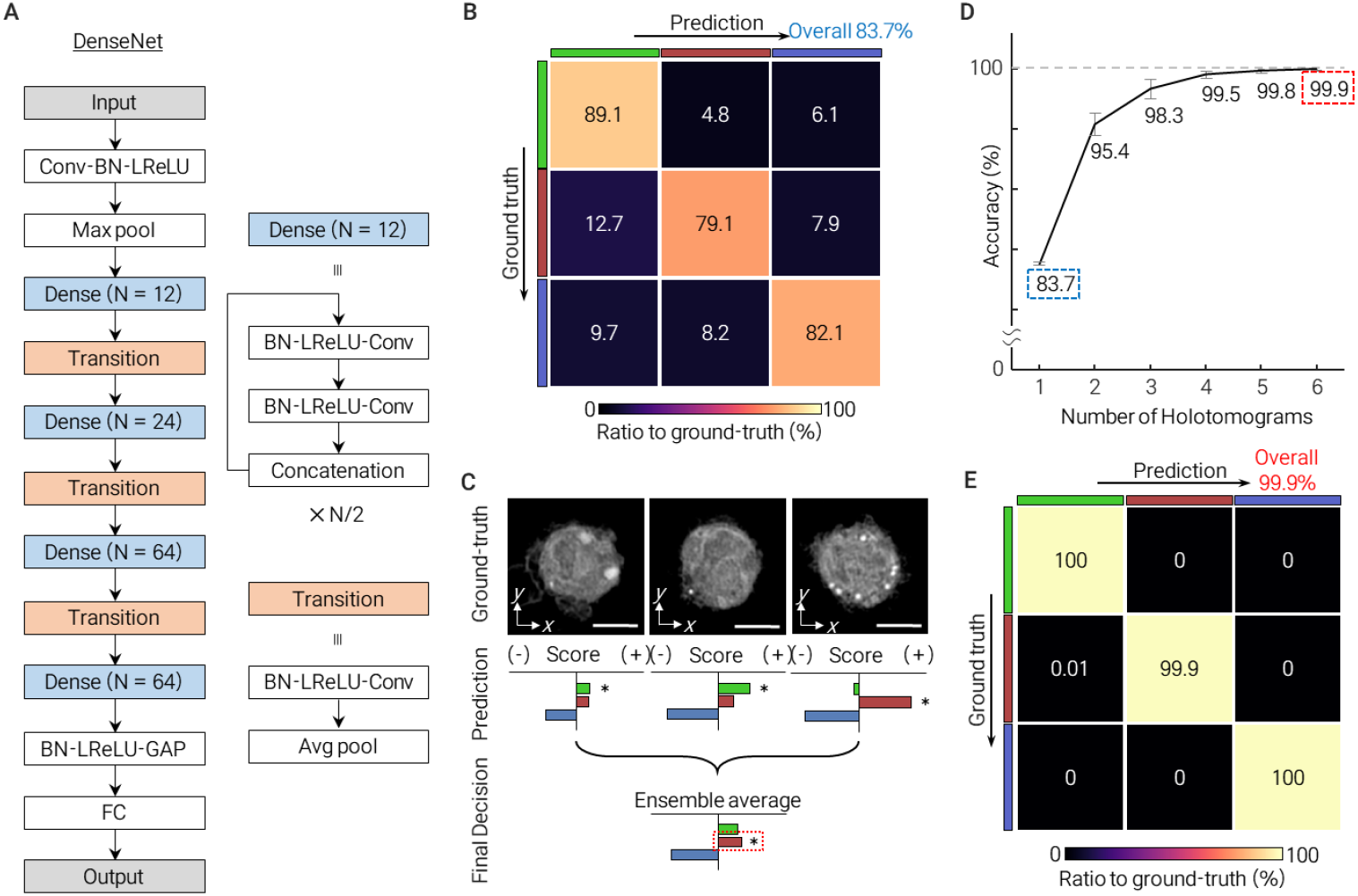
Deep learning-based classification of monocyte immune states using holotomograms. a, Architecture of the 3D convolutional neural network based on DenseNet, comprising multiple dense blocks and transition layers. b, Confusion matrix showing classification performance on single monocyte holotomograms. Overall accuracy = 83.7%. c, Ensemble prediction strategy using multiple monocytes. Three tomograms from the hyper-inflammatory class are shown with individual predictions (asterisks) and final ensemble output (red dashed box). Scale bars = 5 μm. d, Classification accuracy as a function of the number of holotomograms used per prediction. Accuracy improves with ensemble averaging, reaching 99.9% with six cells. e, Confusion matrix based on six monocyte tomograms per prediction. Overall accuracy = 99.9%.

A total of 4,059 monocyte holotomograms were used for training and evaluation. Of these, 407 were reserved for blind testing and 407 for validation, with the remainder used for training. When evaluated on single-cell inputs, the classifier achieved an overall accuracy of 83.7%, with class-wise accuracies of 89.1% (control), 79.4% (hyper-inflammation), and 82.1% (immune suppression) (Fig. 6b). The most common misclassification involved hyper-inflammatory cells being identified as control, likely reflecting their proximity in the early stages of immune activation.

To improve prediction robustness, we employed an ensemble strategy by averaging predictions across multiple monocyte holotomograms. Even when individual predictions were inconsistent, ensemble v oting corrected misclassifications in most cases (Fig. 6c). As the number of monocytes included per prediction increased, accuracy improved substantially—from 83.7% with a single input to 99.9% with six inputs (Fig. 6d). At this level, only one classification error was observed across the entire test set, involving a hyper-inflammatory cell ensemble misidentified as control (Fig. 6e). These results demonstrate the power of combining label-free 3D imaging with deep learning for highly accurate, single-cell–level immune profiling.

## Discussion

In this study, we present a label-free, AI-assisted strategy for immune status classification of individual monocytes using three-dimensional HT and deep learning. HT enabled the non-destructive quantification of intracellular morphological and biophysical features from live, unlabeled cells, which were subsequently used to train a classifier distinguishing control, hyper-inflammatory, and immune-suppressed states. The classifier achieved 83.7% accuracy with single-cell inputs and 99.9% when aggregating predictions from six monocytes. These findings suggest that functional immune phenotypes can be inferred directly from RI-based features without molecular labeling.

Our analysis revealed that LD accumulation strongly correlates with immune dysregulation. Both hyper-inflammatory and immune-suppressed monocytes exhibited significantly increased LD volume and mass, consistent with prior reports of lipid metabolic reprogramming in activated or dysfunctional immune cells ^37^. For instance, LD accumulation has been observed in dendritic cells of cancer patients and macrophages under infection or inflammatory stress ^38^, where it is thought to reflect metabolic reprogramming and immunoregulatory activity^39-42^. These results underscore the relevance of label-free, RI-based lipid detection as a surrogate for immune cell state, and demonstrate the utility of HT in quantifying lipid -rich subcellular structures without staining.

In addition to lipid droplets, we identified a highly discriminative feature—partial protein ratio within the RI range of 1.365–1.370—that likely reflects dense, proteinaceous compartments such as nucleoli or granular cytoplasm. Previous HT studies have associated this RI range with nucleolar structures^27,32^, which are known to undergo remodeling during immune activation or stress. This feature’s predictive power highlights the potential of subcellular RI signatures as functional biomarkers, although further validation with molecular markers is warranted.

While our monocyte population was enriched based on CD14 expression, it likely included a mixture of classical, intermediate, and non-classical subtypes ^43^. Since subtype separation was not performed, the extracted HT features represent averaged phenotypes. Nevertheless, our classifier accurately predicted immune states without explicit subtype distinction, suggesting that immune functional status induces dominant biophysical changes. While previous studies have applied HT to characterize CD8^+^ T cells for sepsis prognosis ^44^, our work demonstrates that similar principles can be extended to monocytes and generalized to multiple immune states. Future studies incorporating CD16-based sorting or transcriptomic validation could refine subtype-specific HT profiles.

Our findings parallel previous work by Sung et al., who used holotomography to classify CD8^+^ T cell morphology in sepsis patients and achieved clinically relevant stratification without labels ^44^. Together, these results suggest that RI-based immune phenotyping can generalize across multiple immune cell types. Although our analysis did not explicitly separate monocyte subtypes, we acknowledge that classical, intermediate, and non - classical subsets—as originally defined by Ziegler-Heitbrock^43^—may exhibit distinct morphological and functional profiles. Future work incorporating subset-specific markers (e.g., CD14/CD16) could refine the interpretation of HT-derived features.

To evaluate model generalizability, we analyzed HT features from 11 donors and observed that inter-status variation exceeded inter-donor variability. This robustness supports the potential of HT-based immune profiling for broader clinical use. However, external validation—such as leave-one-donor-out cross-validation or testing on clinical samples—remains a critical next step to ensure reproducibility.

For clinical translation, reducing imaging and analysis time is essential. Integrating HT with microfluidics - based PBMC enrichment could enable near real-time immune profiling directly from whole blood ^45^. However, current workflows still require improvement in subtype-specific selection, which may be addressed using image-based deep learning trained on paired RI and fluorescence data.

To enhance biological interpretability, future studies should aim to uncover the molecular and functional basis of discriminative HT features. Since RI primarily reflects total protein density, it lacks molecular specificity and cannot distinguish between biomolecular classes or pathways. One promising direction is to train neural networks on correlative datasets that combine 3D holotomograms with molecular readouts—such as cytokine profiles, mHLA-DR expression, or single-cell transcriptomics ^46^. Furthermore, integrating RI imaging with fluorescence-based labels via correlative imaging^28,47,48^, and leveraging such multimodal data in AI training pipelines ^46,49-51^, may enable interpretable, mechanism-informed immune classification. These strategies can bridge the gap between physical phenotype and immunological function, advancing HT from a high -throughput screening tool toward a clinically actionable diagnostic platform.

In conclusion, our study demonstrates that 3D HT combined with AI offers a robust, label-free framework for real-time immune profiling. By capturing functional immune states from single-cell morphology and refractive index features, this approach opens new possibilities for immune monitoring in sepsis, immunotherapy, and precision diagnostics.

## Methods

### Blood collection and monocyte enrichment

This study was approved by the Institutional Review Board (IRB) of KAIST (Approval No. KH2017-004), and all procedures were conducted in accordance with the Declaration of Helsinki. Informed consent was obtained from all participants.

Peripheral blood was collected from healthy donors into EDTA-coated tubes. Mononuclear cells were isolated by density gradient centrifugation using Histopaque®-1077 (Sigma-Aldrich, MO, USA), following dilution of whole blood 1:1 with autoMACS® Rinsing Solution containing bovine serum albumin (Miltenyi Biotec, Germany). A total of 20 mL of diluted blood was carefully layered over 15 mL of Histopaque and centrifuged at 700 g for 30 minutes at room temperature. The mononuclear cell layer was collected and washed twice with buffer.

Monocytes were then enriched using the Pan Monocyte Isolation Kit (Miltenyi Biotec, Germany), following the manufacturer’s protocol. Briefly, PBMCs were incubated with magnetic bead-conjugated antibodies and passed through a magnetic column, collecting the unlabeled monocyte fraction. The resulting monocytes were resuspended in RPMI-1640 medium (Gibco®, USA) supplemented with 10% fetal bovine serum and 1% penicillin/streptomycin (Sigma-Aldrich, USA), and aliquoted for downstream use.

### *In vitro* sepsis model preparation

The in vitro sepsis model was established based on previously reported protocols ^20,52^. To induce the immune suppression (paralysis) state, monocytes were treated with 10 ng/mL LPS for 4 hours at 37 °C in a humidified incubator with 5% CO_2_. Cells were then washed twice with PBS and resuspended in fresh complete medium. After 16 hours, a second LPS stimulation (10 ng/mL, 4 hours) was applied before imaging.

For the hyper-inflammatory condition, monocytes were first cultured for 20 hours in complete medium, followed by a single LPS treatment (10 ng/mL) for 4 hours. Cells were then transferred to imaging dishes at a concentration of 1 × 10^6^ cells per dish. Control monocytes were cultured in complete medium for 24 hours without any stimulation and subsequently transferred to imaging dishes at the same concentration.

### Lipid droplet staining and fluorescence imaging

To validate that high-RI granules observed in holotomograms corresponded to LDs, monocytes were stained with BODIPY™ 493/503 (Invitrogen, Carlsbad, CA, USA), a fluorescent lipid-specific dye. The dye was diluted 1:1000 (v/v) in complete medium, and monocytes were incubated at 37 °C in a humidified 5% CO_2_ atmosphere for 5 minutes. Cells were then washed with PBS to remove excess dye and resuspended in fresh medium.

For fluorescence imaging, 3D image stacks were acquired using wide-field microscopy with a step size of 0.313 μm, followed by 3D deconvolution. The excitation light (centered at 470 nm) intensity and exposure time were manually adjusted to ensure consistent visualization of LD structures across samples.

### HT imaging

The underlying principle of HT is conceptually analogous to inverse scattering, similar to the way computed tomography reconstructs internal structures using X-rays^25,53^. Based on these capabilities, HT has been used to study cell pathophysiology^54,55^, classify cell subtypes^56-58^ and to identify distinct cell pathways ^59^ when combined with artificial intelligence^60^.

Following monocyte isolation and sample preparation, cell suspensions were loaded into customized imaging dishes (Tomodish, Tomocube Inc., Daejeon, Republic of Korea) and covered with a coverslip. After cells settled into a monolayer, three-dimensional RI tomograms of individual monocytes were acquired using a commercial holotomography system(HT-2H, Tomocube Inc.).

The system employs multi-angle Mach–Zehnder interferometry with a 532 nm coherent laser steered by a digital micromirror device (DMD). A total of 49 interferograms were captured under varying illumination angles, and the 3D RI distribution was reconstructed by mapping the measured field data into Fourier space, followed by compensation of the missing cone using 40 iterations of a non-negativity-constrained recovery algorithm. Each tomogram acquisition required approximately 0.5 seconds.

All measurements were performed within 30 minutes per dish to ensure cell viability. After this window, samples were replaced and imaging was resumed. Detailed descriptions of the imaging principles, reconstruction algorithms, and regularization methods are provided in previous reports ^27,34,49,5^.

### Gross morphological features extraction

Morphological features of individual monocytes were quantitatively extracted from RI tomograms acquired using HT. Following segmentation of the region of interest (ROI), we computed geometric descriptors such as cell volume, surface area, aspect ratio, solidity, and sphericity using MATLAB’s built-in region analysis functions, as detailed in the Supplementary Information.

Assuming that the cytoplasm is an aqueous suspension of proteins, the intracellular protein concentration (*C*_*protein*_) can be estimated from the mean RI of the cell (*n*_*cell*_) using the linear relation:

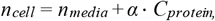

where *n*_*media*_ is the RI of the surrounding medium and *α* = 0.185 mL/g is the RI increment for proteins ^34,35^. The total protein mass was then calculated by multiplying the concentration by the segmented cell volume.

Similarly, the LD concentration and mass were inferred from the segmented LD RI (*n*_*LD*_) using:

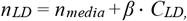

where *C*_*LD*_ is the lipid concentration, and *β* = 0.135 mL/g is the RI increment for lipid components ^35^. The LD mass was computed as the product of lipid concentration and segmented LD volume.

### Deep learning algorithm

Model training was conducted using stochastic gradient descent (SGD) with a mini-batch size of 32. To improve convergence stability, we applied cosine annealing to the learning rate schedule, starting at 0.001 with a cycle length of 32 epochs. The model was optimized using cross -entropy loss and implemented in PyTorch (v1.0.0; https://pytorch.org/).

To reduce overfitting, data augmentation was applied dynamically at each epoch. This included random horizontal cropping, rotation, and Gaussian noise addition. These strategies enhanced generalization by increasing the diversity of the training dataset.

To further boost classification performance, we employed ensemble learning by aggregating predictions from multiple independently trained models. Candidate models were selected based on training and validation accuracy. We evaluated four ensemble strategies—simple averaging, exponential averaging, majority voting, and maximu m probability projection—and optimized additional hyperparameters such as model count, accuracy -based weighting, and output normalization. The ensemble configuration with the highest validation accuracy was used for final prediction.

## Acknowledgments

We thank all the members of KAIST Biomedical Optics Laboratory and Tomocube Inc. for fruitful discussions. We especially thank for department of laboratory medicine of KAIST Clinic Pappalardo Center for kindly helping with blood collection. Images were created with BioRender.com (https://www.biorender.com/) and Power Point (Microsoft, USA).

## Contributions

M.J.L. designed the study with input from Y.K.P. and K.K. Data acquisition was done by M.J.L., J.W.S., Y.S.K., Y.J.C., and J.H.L. M.J.L. and M.S.L. performed data processing and machine learning analysis and D.H.R., and G.K. performed deep-learning analysis. M.J.L. wrote the manuscript, with feedback from M.S.L., G.K., D.H.R., K.K., and Y.K.P.

## Data availability

Data underlying the results presented in this paper are not publicly available at th is time but may be obtained from the authors upon reasonable request.

