## Supplementary Information for "Functional immune state classification of unlabeled live human monocytes using holotomography and machine learning"

[1]*Graduate School of Medical Science and Engineering, Korea Advanced Institute of Science and Technology (KAIST), Daejeon 34141, Republic of Korea*
[2]*Department of Physics, KAIST, Daejeon, Republic of Korea*
[3]*KAIST Institute for Health Science and Technology, KAIST, Daejeon, Republic of Korea*
[4]*Department of Emergency Medicine, CHA Bundang Medical Center, CHA University, Seongnam 13497, Gyeonggi, Republic of Korea*
[5]*Tomocube Inc., Daejeon, Republic of Korea*
[6]*Current affiliation: Division of Pulmonology and Critical Care Medicine, Department of Internal Medicine, College of Medicine, Chungnam National University, Daejeon 34134, Republic of Korea*
[7]*Current affiliation: Institute for Functional Matter and Quantum Technologies, University of Stuttgart, 70569 Stuttgart, Germany*
[8]*Current affiliation: Department of Electrical Engineering & Computer Science, Massachusetts Institute of Technology32 Vassar St, Cambridge, MA 02139, USA*

**Table 1: HT Features**

**Figure 1:**

**Table 1: HT Features**

| Number | Feature name | Meaning | Category |
|---|---|---|---|
| **1** | Cell Volume | Volume of segmented cell | Gross holotomographic features |
| **2** | Cell Surface Area | Surface area of segmented cell | |
| **3** | Surface Volume Ratio | Surface area / volume | |
| **4** | Aspect Ratio | Long axis length/short axis length of segmented cell | |
| **5** | Solidity | Proportion of the voxels in the convex hull that are also in the region | |
| **6** | Sphericity | Sphericity of segmented monocyte | |
| **7** | Protein Mass | Protein content of segmented cell | |
| **8** | Protein Density | Protein density of segmented cell | |
| **9** | Convex Volume | Number of voxels in convex image | |
| **10** | Entropy | Entropy of number of RI distribution | |
| **11** | Mean RI | Mean RI of segmented cell | |
| **12** | RI std | Standard deviation of RI distribution of cell | |
| **13** | Cell Threshold | RI threshold used for cell segmentation | |
| **14** | Equivalent Radius | Diameter of a sphere with the same volume as the cell | |
| **15** | Mean radial distance LD | Mean LD spatial distribution from the centroid | LD-related holotomographic features |
| **16** | Radial distance std LD | standard deviation of spatial LD distribution from the centroid | |
| **17** | LD Volume | Volume of lipid droplets | |
| **18** | LD Mass | Mass of lipid droplets | |
| **19** | LD Number | Number of lipid droplets | |
| **20** | LD Density | Mass density of lipid droplets | |
| **21** | Mean RI per LD | Mean RI of lipid droplets | |
| **22** | Mean volume per LD | Mean single LD volume | |
| **23** | Partial Volume I | Volume occupied by voxel in the RI range | Partial holotomographic features (1.355<RI≤1.36) |
| **24** | Partial Volume Ratio I | The ratio of the partial volume of the RI range to the total volume | |
| **25** | Mean radial distance I | Mean distance of RI distribution from the centroid in the RI range | |
| **26** | Radial distance Std. I | Standard deviation of RI distribution from the centroid in the RI range | |
| **27** | Partial Protein Density I | Mean protein density of segmented regions in the RI range | |
| **28** | Protein Mass I | Protein mass in the RI range | |
| **29** | Partial Protein Ratio I | Ratio of partial protein mass and total protein mass | |
| **30** | Partial Volume II | Volume occupied by voxel in the RI range | Partial holotomographic features (1.36<RI≤1.365) |
| **31** | Partial Volume Ratio II | The ratio of the partial volume of the RI range to the total volume | |
| **32** | Mean radial distance II | Mean distance of RI distribution from the centroid in the RI range | |
| **33** | Radial distance Std. II | Standard deviation of RI distribution from the centroid in the RI range | |
| **34** | Partial Protein Density II | Mean protein density of segmented regions in the RI range | |
| **35** | Protein Mass II | Protein mass in the RI range | |
| **36** | Partial Protein Ratio II | Ratio of partial protein mass and total protein mass | |
| **37** | Partial Volume III | Volume occupied by voxel in the RI range | Partial holotomographic features (1.365<RI≤1.37) |
| **38** | Partial Volume Ratio III | The ratio of the partial volume of the RI range to the total volume | |
| **39** | Mean radial distance III | Mean distance of RI distribution from the centroid in the RI range | |
| **40** | Radial distance Std. III | Standard deviation of RI distribution from the centroid in the RI range | |
| **41** | Partial Protein Density III | Mean protein density of segmented regions in the RI range | |
| **42** | Protein Mass III | Protein mass in the RI range | |
| **43** | Partial Protein Ratio III | Ratio of partial protein mass and total protein mass | |
| **44** | Partial Volume IV | Volume occupied by voxel in the RI range | Partial holotomographic features (1.37<RI≤1.375) |
| **45** | Partial Volume Ratio IV | The ratio of the partial volume of the RI range to the total volume | |
| **46** | Mean radial distance IV | Mean distance of RI distribution from the centroid in the RI range | |
| **47** | Radial distance Std. IV | Standard deviation of RI distribution from the centroid in the RI range | |
| **48** | Partial Protein Density IV | Mean protein density of segmented regions in the RI range | |
| **49** | Protein Mass IV | Protein mass in the RI range | |
| **50** | Partial Protein Ratio IV | Ratio of partial protein mass and total protein mass | |
| **51** | Partial Volume V | Volume occupied by voxel in the RI range | Partial holotomographic features (1.375<RI≤1.38) |
| **52** | Partial Volume Ratio V | The ratio of the partial volume of the RI range to the total volume | |
| **53** | Mean radial distance V | Mean distance of RI distribution from the centroid in the RI range | |
| **54** | Radial distance Std. V | Standard deviation of RI distribution from the centroid in the RI range | |
| **55** | Partial Protein Density V | Mean protein density of segmented regions in the RI range | |
| **56** | Protein Mass V | Protein mass in the RI range | |
| **57** | Partial Protein Ratio V | Ratio of partial protein mass and total protein mass | |
| **58** | Partial Volume VI | Volume occupied by voxel in the RI range | Partial holotomographic features (1.38<RI≤1.385) |
| **59** | Partial Volume Ratio VI | The ratio of the partial volume of the RI range to the total volume | |
| **60** | Mean radial distance VI | Mean distance of RI distribution from the centroid in the RI range | |
| **61** | Radial distance Std. VI | Standard deviation of RI distribution from the centroid in the RI range | |
| **62** | Partial Protein Density VI | Mean protein density of segmented regions in the RI range | |
| **63** | Protein Mass VI | Protein mass in the RI range | |
| **64** | Partial Protein Ratio VI | Ratio of partial protein mass and total protein mass | |
